# Interleukin-6 responses to acute stress are not altered in alcohol use disorder despite elevated baseline inflammation

**DOI:** 10.64898/2026.02.23.707348

**Authors:** Yana Schwarze, Johanna Voges, Sarah Stenger, Janine Stierand, Klaus Junghanns, Oliver Voß, Jennifer Hundt, Frieder Michel Paulus, Sören Krach, Maurice Cabanis, Lena Rademacher

## Abstract

Acute stress activates the immune system, leading to the release of pro-inflammatory cytokines, such as interleukin-6 (IL-6). Chronic alcohol consumption alters the physiological stress systems and is associated with increased chronic inflammation. However, it remains unclear how IL-6 responds to acute stress in individuals with alcohol use disorder (AUD).

Forty patients with AUD during early abstinence and 37 healthy controls (HC) completed two study visits. On one day, an acute stress induction task was performed, and on the other, a non-stressful control task, with the order of tasks being balanced. Plasma IL-6 and C-reactive protein (CRP) were measured as inflammatory markers at baseline and changes in IL-6 were assessed 90 minutes after the experimental manipulation.

Patients with AUD showed significantly elevated baseline IL-6 and CRP compared to HC and the levels correlated positively with the amount of consumed alcohol in patients. IL-6 responses to the stress intervention did not differ between groups. Increases in IL-6 were observed on stress and control days and were larger when samples were collected via an indwelling catheter than with a butterfly needle.

These findings suggest that IL-6 responses to acute stress do not differ between AUD and HC, despite increased baseline inflammation. Furthermore, the results indicate that blood collection methods can influence IL-6 measurements and highlight the importance of methodological considerations.

## 1 Introduction

The body’s response to an acute stressful situation enables individuals to respond adequately to the perceived threat and helps restore homeostasis (Kloet et al., 2005). Stressors trigger a chain of behavioral and physiological responses. The physiological responses are mainly characterized by the activation of the two main stress systems – the hypothalamic-pituitary-adrenal (HPA) axis and the sympathetic nervous system (Kloet et al., 2005). In parallel, the immune system is activated, leading to the release of proinflammatory cytokines, such as interleukin-6 (IL-6) (Slavich and Irwin 2014). In healthy conditions, this inflammatory response is predominantly suppressed by cortisol, released through HPA axis activity (Slavich and Irwin 2014). Importantly, immune activation occurs not only in response to physical stressors or injury, but also in response to psychological stressors, such as experimentally induced psychosocial stress (Steptoe et al. 2007), with IL-6 levels typically peaking around 90 minutes after stressor onset (Marsland et al. 2017).

Chronic activation of physiological stress systems results in alterations of HPA axis and autonomic activity (Chen et al. 2020), as well as a disruption of immune functioning. One factor that causes sustained activation of stress systems is chronic alcohol consumption. Not only has it been shown that chronic alcohol use leads to elevated basal stress system activity, but also blunted responses of the HPA axis to acute stressors (Rachdaoui and Sarkar 2017; Junghanns et al. 2003, 2005; Adinoff et al. 2005). Similarly, previous analyses from our own group showed dampened physiological stress responses in patients with AUD during early abstinence, despite elevated self-reported affective stress experience (Schwarze et al. 2024). In line with chronically elevated stress systems’ activity, higher levels of the inflammatory markers IL-6 (Adams et al. 2020; Moura et al. 2022) and C-reactive protein (CRP) (Costello et al. 2013; Portelli et al. 2019) have been found in patients with AUD in comparison to healthy controls. However, it is yet unclear how chronic alcohol consumption perturbs the inflammatory responses to acute stressful situations, such as the release of IL-6. Given reduced glucocorticoid-mediated suppression, it can be hypothesized that IL-6 might increase more strongly in response to acute stress in these individuals. To our knowledge, only two studies have examined IL-6 responses to acute stress in AUD, both using a personalized stress imagery script and a relaxing, non-physiologically arousing script as a control condition. Fox and colleagues measured cytokine concentrations beginning approximately one hour after insertion of a catheter until 30 minutes (Fox et al. 2020) or 60 minutes (Fox et al. 2017) after the script imagery. The studies reported a reduced IL-6 response of patients with AUD during early abstinence compared to a control group 15 to 30 minutes after the imagery, but no significant differences afterwards. These results contradict a meta-analysis that showed that the effects of stress on IL-6 levels are not significant within the first 30 minutes and increase until approximately 90 minutes after the exposure to the stressor (Marsland et al. 2017). In addition, the findings in both studies were not specific to the stress script imagery, but also occurred for the control condition. Whether or not these deviations from the expected IL-6 response reflect true underlying differences in AUD can also be found in stress paradigms involving social-evaluative elements — which elicit stronger stress responses than imagery paradigms (Dickerson and Kemeny 2004) — or arise from sampling error remains to be clarified.

Importantly, systemic inflammation has also been connected to depressive symptoms: Several studies have found associations between depressive symptoms and circulating IL-6 levels in healthy subjects and patients diagnosed with depression (Mac Giollabhui et al. 2021; Howren et al. 2009), with the connection being driven in particular by somatic symptoms (Duivis et al. 2013). In the context of AUD, however, the relationship appears to be more complex. Neupane et al. (2014) reported that the amount of alcohol intake moderated the relationship between depression severity and IL-6 levels in patients with AUD during early to prolonged abstinence. In particular, they found higher IL-6 levels in AUD patients with major depression compared to non-depressed patients only among individuals consuming lower amounts of alcohol, whereas this difference was absent in patients with higher alcohol use. This could possibly reflect a ceiling effect in heavy drinkers, where alcohol-related systemic inflammation masks depression-related IL-6 elevations, although the exact mechanisms remain unclear.

The central aim of the present study was to investigate differences in the inflammatory response to the acute experience of psychosocial stress, as indicated by IL-6 release, in patients with AUD during early abstinence compared to healthy controls. In addition, we aimed at investigating the association between depressive symptoms and baseline IL-6 in patients with AUD, taking into account the amount of alcohol consumed as a moderator. For this purpose, participants were examined on two separate days, using the Trier Social Stress Test as an acute stress-inducing task on one day and a non-stressful control task on the other (presented in counterbalanced order). Blood samples for IL-6 analysis were collected before and approximately 90 minutes after the end of the intervention to assess inflammatory activity. In addition, questionnaire data on depressive symptoms were obtained. Based on our previous findings of blunted cortisol responses to stress in AUD (Schwarze et al. 2024), we hypothesized that the increase in IL-6 from pre to post stress is higher, due to reduced glucocorticoid-mediated suppression, while baseline levels of inflammatory markers are also higher than in the control group. Furthermore, based on the results of Neupane et al. (2014), we expected a moderating effect of the amount of alcohol consumed on the relationship between baseline IL-6 and depressive symptoms.

## 2 Methods

### 2.1 Sample

Between February 2019 and October 2022 (with an interruption between March 2020 and October 2021 due to the COVID-19 pandemic), 42 individuals diagnosed with moderate or severe AUD according to DSM-5 criteria (meeting 5–11 criteria) and 38 healthy control participants were recruited through the Department of Psychiatry and Psychotherapy in Lübeck and the AMEOS Clinic in Lübeck, Germany. Healthy control subjects were recruited via flyers, letters, and online advertisements. All patients were in the early abstinence phase (between 10 and 40 days) after the acute withdrawal symptoms had subsided and remained in the clinic for the duration of the study. Two patients and one control subject discontinued the study. Thus, the final sample comprised 40 patients and 37 healthy control subjects. Incomplete IL-6 data from two patients and two control participants led to their exclusion from analyses of IL-6 responses to the stress intervention. The two groups were matched for age and gender. All participants provided written informed consent, and the study was approved by the ethics committee of the University of Lübeck (AZ 17-077).

Participants were required to meet the following criteria: age between 18 and 60 years, sufficient knowledge of German, compatibility with the scanner, no neurological disorders or brain injuries, no medications that could have affected the HPA axis or dopamine system in the last two weeks, no acute infections, no vaccinations in the last three weeks, and no chronic inflammatory diseases. Additional inclusion criteria for control participants were: no current or previous psychiatric disorders, no alcohol abuse classified as ≥ 8 points on the Alcohol Abuse Test, no use of drugs except alcohol, nicotine, and occasional cannabis use (less than once per month) in the past year, and no history of regular drug use/dependence except nicotine. Patients additionally had to meet the following criteria: diagnosis of alcohol dependence according to the Diagnostic and Statistical Manual of Mental Disorders, 5th edition (DSM-V), abstinence from alcohol for at least 10 days, no comorbid diagnosis of antisocial or borderline personality disorder, social anxiety disorder, psychosis, acute suicidality, or major depression. Major depression was excluded despite the investigation of depressive symptoms to ensure that AUD was the primary diagnosis and to reduce potential ethical concerns associated with administering a psychologically demanding stress protocol to individuals with heightened vulnerability (Narvaez Linares et al. 2020).

### 2.2 Procedure

Before participating in the study, all participants were informed of the general procedure, but details of the stress-induction protocol were withheld. They were then examined to ensure that they met the inclusion and exclusion criteria and screened for signs of an acute infection within the last two weeks. After the eligibility screening, participants visited the Center for Brain, Behavior, and Metabolism in Lübeck on two different days, with a maximum of 10 days between visits (mean interval: 3.18 days), except for one participant (for details, see Supplement).

The two test days were identical in structure and differed only in the stress protocol, with the order of conditions balanced between participants (see Fig. 1 for an overview). On one day, participants completed a modified version of the Trier Social Stress Test (TSST), a validated procedure for inducing acute psychosocial stress. On the other day, participants performed a control task, following the TSST procedure without inducing psychosocial stress (see below for details of the task). Each test session lasted approximately three hours (8:30– 11:30 a.m. ± 30 minutes). The current study was part of a larger project that also collected MRI data as well as affective and physiological (pulse rate and cortisol) responses, which are not analyzed here, but have been published previously (Schwarze et al. 2024, 2026).

**Figure 1.**
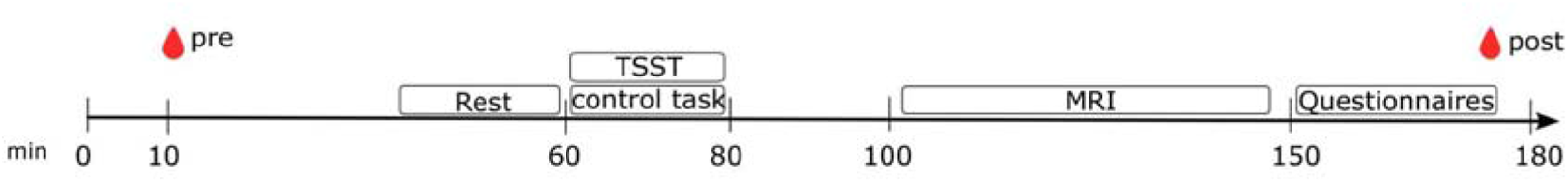
Overview of the experimental design. The two test days followed the same structure and differed only in the stress protocol, with condition order counterbalanced across participants. Blood was collected at the beginning of the test days for baseline values (pre) and 90 minutes after the end of the intervention (post). In addition to the pre and post blood samples, in about half the sample, two additional blood samples were collected at approximately 85 and 150 minutes after start of the test day, but were not further analyzed. The MRI session is shown for visualization purposes only and is not analyzed in the present study. MRI, magnetic resonance imaging; TSST, Trier Social Stress Test

The sessions began with a review of the inclusion and exclusion criteria, followed by the collection of the first blood sample (in participants with catheter insertion, the first blood sample was obtained immediately after placement). Participants were then instructed in the tasks for the MRI session and practiced them. Participants then watched a calming video with landscape scenes to relax and achieve a neutral emotional state. Afterwards, participants completed the German short version of the State-Trait Anxiety Inventory (STAI-S) (Grimm 2009) to assess their current negative affect before performing either the TSST or the control task, followed by another STAI-S assessment. In addition, participants’ resting heart rates were measured during the calming video (baseline) and during the anticipation and task phases of the TSST/control task using a pulse oximeter (for details, see Schwarze et al. 2024). About twenty minutes after the TSST/control task, a one-hour MRI session followed (see supplement for details). Finally, participants completed several questionnaires (see details below), and another blood sample was taken 90 minutes after the end of the intervention for IL-6 analysis. Participants were debriefed after the stress induction session. On both test days, saliva samples were collected at five time points from the start of the test day to approximately 80 minutes after the stress induction/control task for cortisol analysis (for details, see Schwarze et al. 2024). Please note that after the interruption due to the COVID-19 pandemic, there were some procedural changes, for example, a SARS-CoV-19 rapid test was conducted upon arrival of the participant. In addition, for organizational reasons, blood was no longer collected via an indwelling venous catheter (n = 48; 21 AUD / 27 HC) but via venipuncture (n = 29; 19 AUD / 10 HC), and the number of blood samples taken was reduced from four time points to two. Due to the change in the blood sampling method, which may have an impact on the results, the sampling method was included as a covariate in all analyses evaluating the time course of IL-6.

### 2.3 Stress induction

The Trier Social Stress Test (Kirschbaum et al. 1993) was used to induce acute psychosocial stress in participants, which is a reliable and widely used instrument for investigating acute psychosocial stress under experimental conditions (Allen et al. 2017). The TSST is a standardized protocol that includes a 10-minute anticipation phase followed by a 10-minute test phase. According to the protocol, participants are first familiarized with the tasks they have to complete, including a five-minute free speech and a mental arithmetic task in front of a jury. The jury consists of three people who are briefly introduced to the participant in a second room to increase psychosocial stress. For the anticipation phase, participants return to the first room and are given pen and paper to prepare for the upcoming task while the experimenter waits outside. After ten minutes, the experimenter brings the participant back to the jury room, and the test phase begins, consisting of the five-minute speech, in which the participant must convince the jury that they are the perfect candidate for a fictional job vacancy, and five minutes of mental arithmetic, in which the participant must serially subtract 13 from the number 1022. In the current study, the TSST protocol was followed with only minor deviations: the experimenter did not leave the room during the test phase but was part of the jury, and no microphone was used. As a cover story, participants were told that the task was to test their verbal and mathematical abilities. Five patients were unable to complete the entire task phase of the TSST because they felt too stressed. As the goal of the intervention, stress induction, was clearly achieved in all these cases, we decided to continue with the rest of the procedure. In addition to the stress induction procedure, a control session (similar to (Het et al. 2009) was conducted on a separate day, in which participants only read written texts (instead of free speech as in (Het et al. 2009) and performed simple mental arithmetic tasks in the absence of a jury. This was very similar to the stress induction procedure (both took place in the same rooms, participants were in a standing position during the task phase, and both included verbal and mathematical tasks), but was not intended to cause psychosocial stress related to performance evaluation. Here, participants were informed that the purpose of the tasks was to assess the physiological effects of mental effort by measuring their pulse rate as part of the cover story (when the task was performed on the first day of testing).

### 2.4 Blood samples

Blood samples collected at the beginning of the test day included EDTA tubes for the analysis of IL-6 and, on the first day of testing, for CRP and a complete blood count. At the end of the test day, approximately 90 minutes after the experimental manipulation, another tube for the analysis of IL-6 was taken. After collection, IL-6 blood samples were centrifuged at 1000g for 15 minutes, blood plasma was collected and stored at −80 degrees Celsius until further analysis. The complete blood count and CRP were determined at an external laboratory (Laborärztliche Gemeinschaftspraxis Lübeck). After study completion, IL-6 was measured in duplicates using high-sensitivity enzyme-linked immunosorbent assays (Human IL-6 Quantikine HS SixPak). Intra- and inter-assay coefficients of variation were below 10.57% and 12.89% respectively with a lower limit of detection of 0.313 pg/ml.

### 2.5 Depressive Symptoms

The Beck Depression Inventory (BDI-II) (Hautzinger et al. 2006) was used to assess depressive symptoms in the last week. Since alcohol withdrawal can be accompanied by numerous somatic complaints which can affect scoring in the inventory (Beck et al. 2013) and IL-6 may be differently associated with cognitive and somatic symptoms (Duivis et al. 2013), we analyzed the cognitive and somatic-affective factor of the BDI-II separately (Keller et al. 2008).

### 2.6 Statistical analyses

IL-6 and CRP concentrations were log-transformed prior to analysis to improve normality and homogeneity of variance. To assess baseline differences in IL-6 between the groups, two-tailed Welch’s two-sample t tests were computed for mean baseline levels of both test days.

Separate linear regression models were calculated to examine positive associations with baseline IL-6 and CRP as outcomes and age, BMI, and depressive symptoms (BDI-II cognitive factor and BDI-II somatic-affective factor) as predictors. In each model, group (HC vs. AUD) and the interaction term between the respective predictor and group were included to test whether associations differed between groups. To examine associations between baseline IL-6 or CRP and the amount of alcohol consumed, separate linear regression analyses were conducted within the AUD group only. In order to analyse whether the relationship between baseline IL-6 and depressive symptoms could be moderated by daily alcohol intake in the AUD group, another linear regression was conducted, which included depressive-symptom scores, daily alcohol consumption, and their interaction term. All predictors were mean-centered prior to analysis. To analyze the effects of stress induction on circulating IL-6 levels, a repeated-measures analysis of variance with *group* (AUD vs. HC) as between-subject factor, *condition* (stress vs. control task) and *timepoint* (pre vs. post intervention) as within-subject factors was calculated. *Sampling method* (catheter vs. butterfly) was included as a covariate to explore whether the blood sampling procedures could have impacted the IL-6 response.

Standardized residuals from all regression models were inspected using Q–Q plots, histograms, and Shapiro–Wilk tests, indicating that there were no statistically significant deviations from normal distribution. All data was analyzed using R version 4.3.2.

## 3 Results

### 3.1 Sample characteristics and baseline inflammatory parameters

Information on demographics, clinical characteristics, and blood parameters is reported in Table 1 and Supplementary Table S1. Baseline levels of IL-6 were averaged across both test days as there was no significant difference between the stress and control days (for details, see Supplement).

**Table 1.** Sample characteristics.

| | AUD ($N = 40$)<sup>a</sup> | HC ($N = 37$)<sup>b</sup> | Cohen's $d$ [95% CI] | $t$-statistic | $p$-value |
| --- | --- | --- | --- | --- | --- |
| Gender (% male) | 95 | 94.6 |  |  |  |
| Age (years) | 42.23 ± 9.15 | 43.38 ± 9.68 | | | |
| BMI | 24.84 ± 3.56 | 26.86 ± 4.95 | -0.47 [-0.93, -0.01] | -2.05 | 0.04 |
| Abstinence (days) | 14.31 ± 6.30 | | | | |
| Daily alcohol (g) | 289.13 ± 193.12 | 2.69 ± 3.39 | 2.08 [1.51, 2.65] | 9.14 | < 0.001 |
| Fulfilled DSM-5 criteria | 8.50 ± 2.14 | | | | |
| Years of chronic alcohol consumption | 9.42 ± 11.02 | | | | |
| Baseline STAI-S | 23.74 ± 8.61 | 23.03 ± 7.55 | 0.09 [-0.37, 0.54] | 0.39 | 0.698 |
| BDI-II cognitive factor | 4.60 ± 3.58 | 1.25 ± 1.34 | 1.22 [0.72, 1.71] | 5.51 | < 0.001 |
| BDI-II somatic-affective factor | 7.00 ± 6.28 | 3.21 ± 3.38 | 0.74 [0.27, 1.21] | 3.32 | 0.002 |
| BDI-II total score | 11.60 ± 9.02 | 4.46 ± 4.23 | 1.00 [0.51, 1.48] | 4.49 | < 0.001 |
| Baseline IL-6 (pg/ml) | 2.23 ± 1.87 | 1.16 ± 0.88 | 0.79 [0.32, 1.26] | 3.51 | < 0.001 |
| CRP (mg/l) | 3.75 ± 4.23 | 1.70 ± 1.63 | 0.55 [0.08, 1.01] | 2.42 | 0.02 |
| Leukocytes (10<sup>9</sup>/l) | 7.36 ± 2.20 | 5.42 ± 1.26 | 1.07 [0.59, 1.56] | 4.80 | < 0.001 |
| Thrombocytes (10<sup>9</sup>/l) | 327.75 ± 96.21 | 236.89 ± 52.31 | 1.20 [0.70, 1.69] | 5.20 | < 0.001 |

*Note.* AUD, alcohol use disorder; BDI-II, Beck Depression Inventory II; BMI, body mass index; CRP, C-reactive protein; HC, healthy controls; IL6, interleukin-6; STAI-S, State-Trait Anxiety Inventory (state).

Baseline IL-6, CRP, leukocytes, and thrombocytes were significantly elevated in patients with AUD in comparison to healthy controls (Table 1). IL-6 was significantly and positively correlated with CRP (*r* = 0.70, *p* < 0.001), leukocytes (*r* = 0.52, *p* < 0.001), and thrombocytes (*r* = 0.36, *p* = 0.001).

Values are given as mean and standard deviation if not specified otherwise. For the comparison of groups, Welch t-tests were computed. Baseline levels of IL-6 and STAI-S were averaged across both test days. For one AUD patient and one control participant, the baseline IL-6 value was unavailable on one test day; therefore, the mean baseline IL-6 estimate for these individuals is based on the single available value. ^a^For one AUD participant, the exact duration of abstinence (10–40 days) and the fulfilled DSM-5 criteria is unknown; data on the amount of daily alcohol consumption before withdrawal and years of chronic alcohol consumption are missing for two AUD participants, resulting in a sample size of *N* = 39, N = 39, N = 38, and *N* = 38 for these variables, respectively. ^b^The BDI score is missing in the case of one control participant. Therefore, the sample size is reduced to N=36.

### 3.2 Baseline IL-6 correlates with age and BMI, and in patients with AUD with the daily amount of consumed alcohol

As reported in the literature, baseline IL-6 was positively associated with age (Ferrucci et al. 2005; Capuron et al. 2011) and BMI (Bowker et al. 2020) (see Table 2, Fig. 2). In contrast, cognitive and somatic-affective depressive symptoms were not significantly associated with IL-6 levels. Within the AUD group only, alcohol consumption was positively associated with IL-6. In addition, significant main effects of group were found for BMI, age, and BDI-II somatic-affective factor, indicating overall differences between the groups. However, none of the interactions between predictor and group reached statistical significance, indicating that associations between IL-6 and BMI, age, and depressive symptoms did not significantly differ between HC and AUD participants. Similar patterns were observed for CRP (supplementary Table S2).

**Table 2.** Linear regression analyses of associations with baseline IL-6.

|  | BMI | Age | BDI-II cognitive factor | BDI-II somatic-affective factor | Daily alcohol |
| --- | --- | --- | --- | --- | --- |
| Variable | **0.058 (0.018),**<br>**p = 0.001** | **0.014 (0.008),**<br>**p = 0.046** | 0.048 (0.047),<br>p = 0.151 | 0.005 (0.020),<br>p = 0.405 | **0.001 (0.0007),**<br>**p = 0.034** |
| Group | **0.666 (0.151),**<br>**p < 0.001** | **0.564 (0.156),**<br>**p < 0.001** | 0.361 (0.228),<br>p = 0.059 | **0.511 (0.177),**<br>**p = 0.003** | - |
| Var x group | -0.030 (0.036),<br>p = 0.208 | -0.016 (0.017),<br>p = 0.166 | -0.114 (0.093),<br>p = 0.113 | -0.041 (0.039),<br>p = 0.148 | - |
| R<sup>2</sup> | 0.274 | 0.183 | 0.149 | 0.147 | 0.090 |
| adj. R<sup>2</sup> | 0.244 | 0.149 | 0.114 | 0.112 | 0.064 |
| N | 77 | 77 | 76 | 76 | 38 (AUD only) |

*Note.* Reported p-values are one-sided based on a priori directional hypotheses. Bold values indicate statistical significance at $\alpha = 0.05$ .

**Figure 2.**
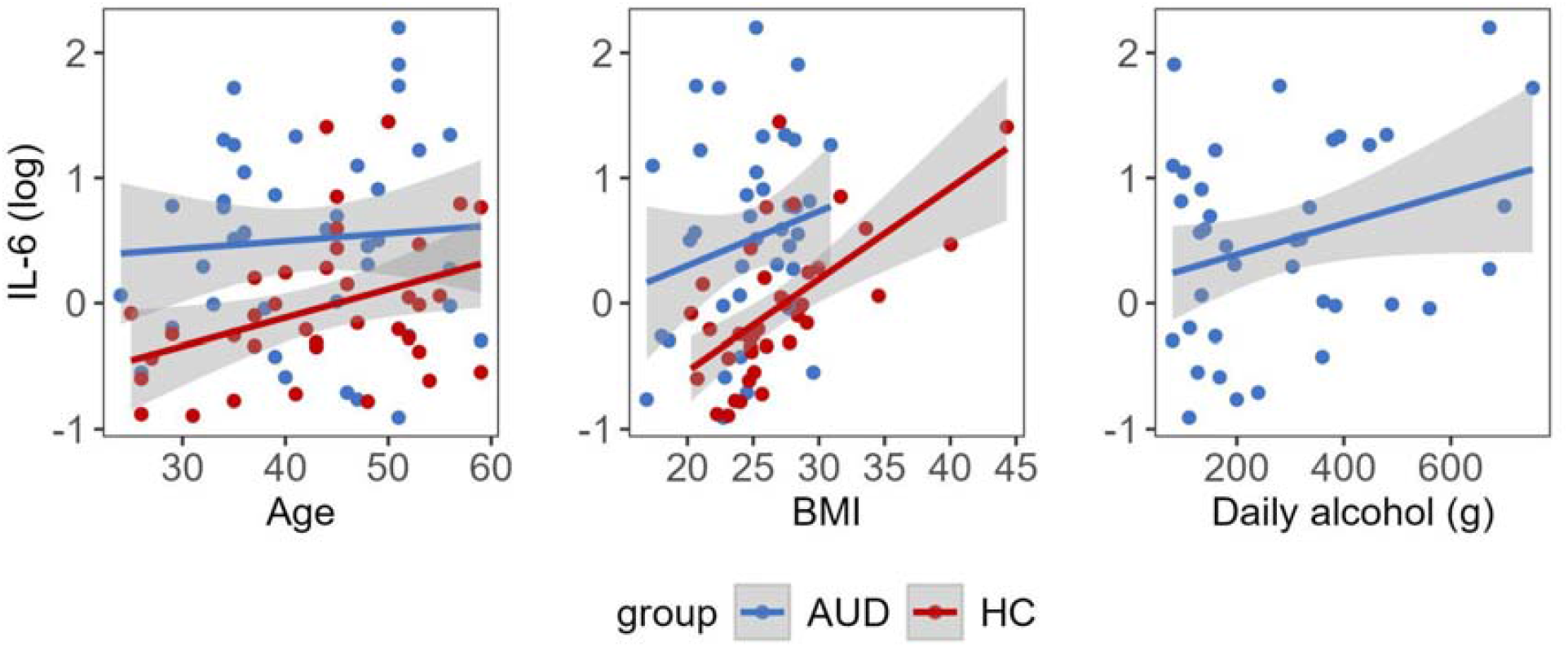
Associations of baseline interleukin-6 (IL-6) with age and Body Mass Index (BMI) and with the amount of alcohol consumed. Scatterplots show the data of each group in case of age and BMI or only patients with alcohol use disorder (AUD) in the case of consumed alcohol overlaid with a regression line and respective confidence resulting from a multiple regression analysis controlling for group differences in IL-6 (log-transformed) concentration. Only main effects were statistically significant (p < 0.05, one-sided).

### 3.3 No moderating effect of daily alcohol intake on the association of IL-6 with depressive symptoms in patients with AUD

To test whether daily alcohol intake moderated the relationship between baseline IL-6 and depressive symptoms in the AUD group, we examined the interaction between depressive symptoms and daily alcohol consumption. For both BDI-II factors, the interaction term was not significant (BDI-II cognitive factor: *t* = −0.07, *p* = 0.947; BDI-II somatic-affective factor: *t* = −0.54, *p* = 0.59), indicating no evidence of a moderating effect.

### 3.4 No alterations in stress-induced IL-6 responses in AUD

Stress was successfully induced on the stress day, but not on the control day, as determined by pulse rate, cortisol levels, and affect ratings (for details, see Schwarze et al. 2024). As expected, the repeated-measures ANOVA on the effects of stress on IL-6 levels revealed generally elevated IL-6 levels in patients with AUD (main effect of *group*: *F*(1, 70) = 11.36, *p* < 0.001). However, the hypothesized effect of stronger IL-6 increases in the AUD group under stress could not be detected (three-way interaction of *group* x *condition* x *timepoint*: *F*(1, 70) = 0.01, *p* = 0.911) (Fig. 3). IL-6 levels were higher after the intervention than at baseline across both test days (main effect of *timepoint*: *F*(1, 70) = 19.34, *p* < 0.001), suggesting an increase independent of the experimental intervention. In addition, IL-6 levels increased more strongly when samples were collected via an indwelling catheter than via a butterfly needle (*sampling method x timepoint interaction*: *F*(1, 70) = 5.42, *p* = 0.023). No other main or interaction effects were found (see Supplementary Table S3 for details).

**Figure 3.**
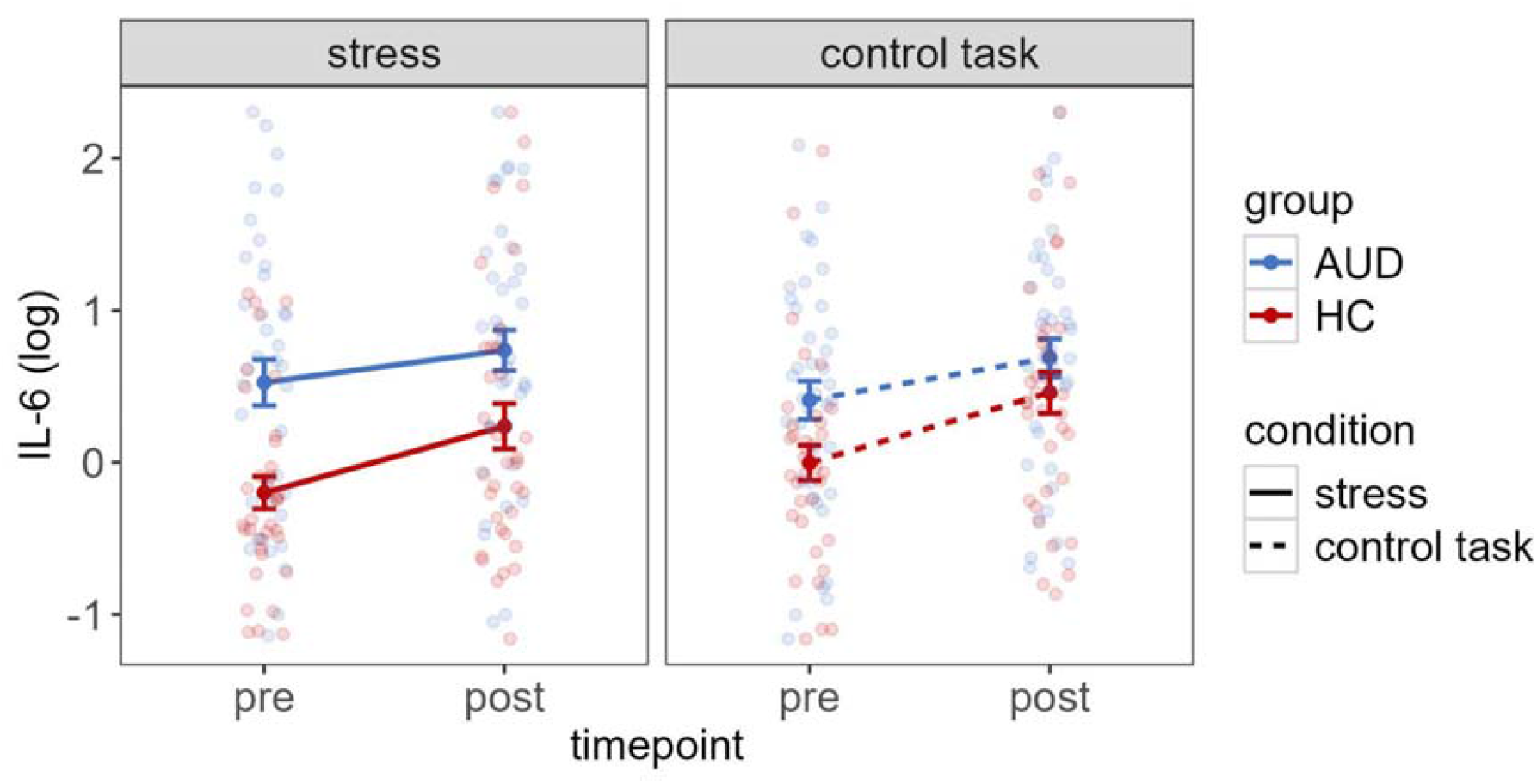
Interleukin-6 (IL-6) responses on stress and control days in patients with alcohol use disorder (AUD) and healthy controls (HC). IL-6 concentration (log-transformed) is displayed at baseline and approximately 90 minutes after the experimental manipulation. Error bars represent standard error of the mean.

## 4 Discussion

The current study investigated the effects of acute psychosocial stress on IL-6 levels in patients with AUD during early abstinence, compared to healthy controls. Baseline IL-6 was significantly higher in the AUD group, but we found no evidence for altered IL-6 responses to acute stress compared to healthy controls.

In line with previous findings, patients with AUD exhibited elevated baseline IL-6 (Moura et al. 2022; Adams et al. 2020) and CRP (Gürbüzer and Özcan Tozoğlu 2024; Xu et al. 2020; Di Gennaro et al. 2007) levels compared to healthy controls. IL-6 and CRP are recognized markers for systemic inflammation (Fonseca et al. 2009; Pay and Shaw 2019) and likely reflect damage from prior heavy alcohol use: Chronic alcohol exposure activates toll-like receptor systems that increase intestinal permeability and facilitate translocation of microbial products into the circulation. These processes potentiate liver inflammation and promote IL-6 release and downstream hepatic CRP synthesis, while peripheral cytokines and chemokines signal to the brain and activate central microglia and astrocytes to release central nervous system cytokines (Neupane 2016). Baseline IL-6 and CRP were positively associated with both age and BMI. This is in line with previous findings (Ferrucci et al. 2005; Capuron et al. 2011; Choi et al. 2013) and reflects the increase of pro-inflammatory cytokine levels with aging referred to as inflammaging (Franceschi et al. 2006), and the release of inflammatory cytokines from adipose tissue (Brooks et al. 2010). Both the elevated baseline CRP and IL-6 in AUD, as well as the association of these markers with age and BMI, validate the measurement of inflammatory processes in the present study.

Patients with AUD showed significant associations of baseline IL-6 or CRP with the amount of alcohol consumed per day, which is consistent with reports linking IL-6 to breath alcohol concentration in AUD (Heberlein et al. 2014) and IL-6 and CRP to the average alcohol consumption (Hsu et al. 2023; Battista et al. 2023). These findings suggest that chronic alcohol consumption contributes to systemic inflammation in a dose-dependent manner. We did not find evidence for altered stress-related IL-6 responses in individuals with AUD during early abstinence compared to HC, although we observed stress-specific group differences for other stress parameters (cortisol, pulse rate, affect) in previous analyses (for details, see Schwarze et al. 2024). This is in line with Fox et al. (2020, 2017), who also did not find stress-specific alterations in IL-6 responses in AUD. The authors reported blunted IL-6 responses in AUD patients within the first 30 minutes after both a stress and a control intervention, but no significant group differences afterwards. In the present study, IL-6 responses were comparable in both groups, with similar increases from baseline to 90 minutes after intervention, occurring not only in the stress condition, but also in the control condition. Although the control task did not elicit changes in cortisol, heart rate, or negative affect in healthy controls, IL-6 levels still increased. This contrasts with the anticipated stress-specific increase in IL-6 reported in the meta-analysis by Marsland et al. (2017). It is worth noting, however, that the studies included in this meta-analysis mostly assessed stress without a non-stressful control task, thereby limiting conclusions about whether the effects are specific to stress, as induced by the task, or relate to other confounding variables. To elaborate this point, when focusing exclusively on the stress condition, our data would also show significant IL-6 increases, aligning with the stress-related cytokine responses reported by Marsland et al. (2017). Of the few previous studies that included a control task, results were heterogeneous, with some studies reporting IL-6 increases specific to the stress task (Brydon et al. 2005; Steptoe et al. 2001; Kuebler et al. 2015), while others observed IL-6 increases in both stress and control conditions (Lutgendorf et al. 2004; Yamakawa et al. 2009; Dugué et al. 1993; Fox et al. 2020, 2017). These mixed results suggest that factors unrelated to the stress manipulation itself may also influence IL-6 responses. One possible explanation could be that the context of the study itself can also trigger a stress response (Gossett et al. 2018). However, another possibility is that the physiological response to blood sampling may also play a role here. Several studies comparing cytokine trajectories across sampling methods have shown that indwelling catheters can induce a local inflammatory response, particularly for IL-6, which is not detectable when using venipuncture (Gudmundsson et al. 1997; Chabot et al. 2018; Haack et al. 2002, 2000). The use of an indwelling catheter is the most common approach in studies assessing inflammatory responses to stress, as it minimizes participant burden and allows for repeated sampling. Many studies, including those by Fox and colleagues (2020, 2017), included a 20–30 minute rest period between catheter insertion and the first baseline sample. However, given that catheter insertion itself elicits a local inflammatory reaction that increases over time, its use may influence the measurement of inflammatory parameters and thereby bias the interpretation of cytokine trajectories. It should be noted that, for organizational reasons, we had to change the blood collection method during the study and found that IL-6 levels increased more strongly (on both study days) in participants whose samples were taken via an indwelling catheter than in participants whose samples were taken via a butterfly needle. Thus, these increases in IL-6 may not be uniquely attributable to the stress intervention, but may partly reflect physiological or psychological responses associated with the presence of the intravenous catheter. To address this potential confound, blood sampling method was included as a covariate in our analysis. Nevertheless, blood sampling method remains an important consideration for all studies on acute stress involving analyses of inflammatory parameters, particularly of IL-6.

We did not find any associations between IL-6 and depression scores in the AUD group and only a statistical trend toward a positive correlation between IL-6 and the BDI-II cognitive factor in the HC group. This contrasts with meta-analyses that report positive correlations between depressive symptom severity and IL-6 in depressed and healthy samples, with stronger effects for clinically depressed patients than community-based samples (Mac Giollabhui et al. 2021; Howren et al. 2009). Also in patients with AUD, a positive association between IL-6 and depression scores has been reported (Martinez et al. 2018). However, a study by Neupane et al. (2014) showed that this relationship can be moderated by the amount of alcohol consumed: The authors demonstrated that among individuals with AUD and lower alcohol consumption, those with a comorbid major depression exhibited elevated IL-6 levels compared with their non-depressed counterparts, whereas among individuals with heavy alcohol consumption there was no difference in IL-6 levels between participants with and without major depression. There was no significant moderating effect of the amount of alcohol consumed on the relationship between IL-6 and depressive symptoms in our sample. One potential explanation for the discrepancy between our results and those of previous studies could be differences in symptom severity. Because major depression was an exclusion criterion in the present study, depressive symptoms were likely lower than in the sample of Martinez and colleagues, who included individuals from a study site specialized in the treatment of persons with depression or in the study by Neupane and colleagues, who specifically recruited patients with major depression. It is possible that a relationship between IL-6 and depressive symptoms only becomes apparent when individuals with severe symptoms are also included. Thus, the restricted range of depressive symptom severity in our sample, together with a rather small sample size, may have constrained the statistical power needed to identify a moderating influence of alcohol consumption.

Several limitations should be considered when interpreting the results. First, our sample consisted predominantly of male participants. It remains unclear whether the findings generalize to females, as sex differences in IL-6 responses to stress have been reported (Edwards et al. 2006; Endrighi et al. 2016). However, re-running all analyses after excluding the few female participants did not yield any significant changes (for details, see Supplement). Second, the groups differed on several variables such as cigarette and cannabis smoking, stress related with the inpatient treatment, or potentially altered health status. Smoking, for example, has been associated with variations in IL-6 concentrations (Elisia et al. 2020). However, given the high prevalence of smoking among patients with AUD, disentangling the specific effects of smoking from those of AUD remains challenging. Third, the current study focused on individuals with AUD during early abstinence. As reported in a meta-analysis by Adams and colleagues (2020), larger differences in cytokine concentrations were observed during active drinking and withdrawal compared to early or prolonged abstinence. Additionally, rigorous exclusion criteria were applied to reduce comorbidity and polysubstance use. Consequently, the findings may not generalize to other phases of AUD or to broader, more heterogeneous AUD populations. Fourth, as part of a larger project, participants underwent an MRI scan, which may have introduced additional stress influencing IL-6 responses. However, as this procedure was identical across all participants, it is unlikely to account for the absence of group differences in the IL-6 response to stress. Finally, the relatively small sample size may have limited the statistical power for some sub-analyses, and these results should therefore be interpreted with caution.

In conclusion, we did not find evidence for altered IL-6 responses to acute stress in the early abstinent AUD group compared to HC, despite increased baseline levels of IL-6 and CRP. IL-6 increases were observed on stress and control days, with larger changes occurring when samples were collected via an indwelling catheter than via a butterfly needle. These findings suggest that factors inherent to experimental procedures can influence cytokine patterns, underscoring the need for control tasks and careful methodological consideration when measuring and interpreting inflammatory responses in stress research. Additionally, our results demonstrate that the typical associations between inflammatory markers and physiological factors observed in healthy controls were absent in the AUD group, while they correlated with the amount of alcohol consumed. This suggests that chronic heavy alcohol consumption affects inflammatory processes to such an extent that the interaction with other physiological processes is altered.

## Acknowledgements

We thank Alexander Schröder, Julie Forster, Emil Zilikens, and Jovana Lehmann-Grube for their help in data collection as well as Birte Hell and Nadine Merg for their help with laboratory analyses.

## Data Availability

Data and code of this study are openly available at: https://osf.io/d7bh2/

