## Supplementary material for "Interleukin-6 responses to acute stress are not altered in alcohol use disorder despite elevated baseline inflammation"

### **Supplementary methods**

#### **MRI sessions**

The present analysis is part of a larger project, which aimed to examine affective, physiological, and neural responses to acute psychosocial stress in AUD, including two MRI experiments. The first experiment was a well-established social adaptation (Spreckelmeyer et al. 2009) of the monetary incentive delay task by Knutson et al. (2000) to examine participants' neural responses to anticipated rewards. During the task, participants respond as quickly as possible to a target stimulus after visual cues indicate a potential reward, with rapid responses yielding either social rewards (smiling faces) or monetary rewards (small amounts of money). The second experiment involved a novel paradigm designed to examine neural and behavioural responses to images of social situations. The participants were presented with images depicting both social and non-social scenes, featuring varying numbers of people. They were then invited to indicate their level of enjoyment of the images.

#### **Information on the participant with the long interval between test days**

One control participant was unable to return for the second test day within the 10-day interval because of illness. Due to his work schedule, he was only able to return for the second test day after 48 days.

#### **Analysis software**

We used R version 4.3.2 (R Core Team 2023) and the following R packages: afex v. 1.3.0 (Singmann et al. 2023), arsenal v. 3.6.3 (Heinzen et al. 2021), cocor v. 1.1.4 (Diedenhofen and Musch 2015), effsize v. 0.8.1 (Torchiano 2025), here v. 1.0.1 (Müller 2020), irr v. 0.84.1(Gamer et al. 2019), kableExtra v. 1.4.0 (Zhu 2024), lme4 v. 1.1.35.1 (Bates et al. 2015), lpSolve v. 5.6.23 (Csárdi and Berkelaar 2025), Matrix v. 1.6.5 (Bates et al. 2024), patchwork v. 1.3.0 (Pedersen 2025), rstatix v. 0.7.2 (Kassambara 2023), smplot2 v. 0.1.0 (Min 2024), tidyverse v. 2.0.0 (Wickham et al. 2019).

### **Supplementary results**

#### **Substance use**

Table S1. Substance and nicotine use in the AUD and control groups.

| Drug and nicotine use | Frecueny | AUD (n) | HC (n) |
| --- | --- | --- | --- |
| Nicotine | Current | 37 | 4 |
|  | Former | 1 | 6 |
|  | Unknown | 1 | 0 |
| Cannabis | Daily | 4 | 0 |
|  | 1-3x/week | 3 | 0 |
|  | Sporadically | 2 | 1 |
|  | Frequency unknown | 2 | 0 |
| Cocaine | Sporadically | 8 | 0 |
| Amphetamines | Sporadically | 3 | 0 |

#### **Baseline inflammatory parameters**

Baseline levels of IL-6 did not significantly differ between the stress and control days (*t*(74) = 0.22, *p* = 0.83, mean difference = –0.02, 95% CI [–0.14, 0.17]) across both groups. This was also the case when examining the AUD group (*t*(38) = –1.37, *p* = 0.18, mean difference = –0.14, 95% CI [–0.34, 0.07]), and the healthy control group (*t*(35) = 1.62, *p* = 0.11, mean difference = 0.18, 95% CI [–0.05, 0.41]) separately. Intraclass correlations (ICC) indicated moderate to high consistency across test days (ICC = .66 for the full sample, 0.72 for the AUD group, and 0.44 for the HC group). The lower ICC observed in healthy controls likely reflects the generally low IL-6 levels in this group.

#### **Supplementary correlations**

Table S2. Linear regression analyses of associations with CRP

|  | BMI | Age | BDI-II cognitive  factor | BDI-II somatic-  affective factor | Daily alcohol |
| --- | --- | --- | --- | --- | --- |
| Variable | **0.120 (0.030),**  **p < 0.001** | 0.014 (0.008),  p = 0.215 | 0.095 (0.078),  p = 0.114 | -0.017 (0.023), p = 0.701 | **0.002 (0.004),**  **p = 0.024** |
| Group | **0.883 (0.249),**  **p < 0.001** | **0.654 (0.266),**  **p = 0.008** | 0.278 (0.383),  p = 0.235 | **0.667 (0.298),**  **p = 0.014** |  |
| Var x group | -0.012 (0.060),  p = 0.418 | -0.039 (0.029),  p = 0.090 | -0.091 (0.157),  p = 0.281 | -0.041 (0.066),  p = 0.267 |  |
| R² | 0.256 | 0.102 | 0.087 | 0.085 | 0.105 |
| adj. R² | 0.225 | 0.065 | 0.049 | 0.047 | 0.080 |
| N | 77 | 77 | 76 | 76 | 38 (AUD only) |

*Note.* Reported p-values are one-sided based on a priori directional hypotheses. Bold values indicate statistical significance at α = 0.05.

#### **ANOVA results on IL-6 responses**

Table S3. Results of the ANOVA on IL-6 responses to acute stress.

|  | *df* | *F* | *η_p_* | *p-*value |
| --- | --- | --- | --- | --- |
| group | 70 | 11.35 | 0.14 | < 0.001 |
| condition | 70 | 0.55 | 0.01 | 0.460 |
| timepoint | 70 | 19.34 | 0.22 | < 0.001 |
| sampling method | 70 | 2.36 | 0.03 | 0.129 |
| group x condition | 70 | 3.70 | 0.05 | 0.059 |
| group x timepoint | 70 | 1.07 | 0.02 | 0.304 |
| condition x timepoint | 70 | 0.39 | 0.01 | 0.537 |
| sampling method x condition | 70 | 0.25 | < 0.01 | 0.616 |
| sampling method x timepoint | 70 | 5.42 | 0.07 | 0.023 |
| group x condition x timepoint | 70 | 0.01 | < 0.01 | 0.911 |
| sampling method x condition x timepoint | 70 | 0.35 | <0.01 | 0.553 |

*Note*. For the ANOVA on IL-6 responses to acute stress, data of two AUD patients and two control participants are missing, resulting in a sample size of N = 38 in the AUD group and N = 36 in the HC group.

#### **Analyses excluding female participants**

To ensure that the results were not biased by the small number of female participants, all analyses were repeated after removing them from the analysis. Rerunning the analysis without any female participants had no substantial impact on the results. Baseline IL-6 and CRP remained significantly elevated in patients with AUD, and inflammatory markers continued to be associated with age and BMI and with alcohol consumption in the AUD group. The amount of alcohol consumed again did not moderate the relationship between IL-6 and depressive symptoms. The ANOVA examining stress effects on IL-6 levels revealed main effects of group and timepoint, as well as a sampling method × timepoint interaction.
